# Early life adversity has sex-dependent effects on survival across the lifespan in rhesus macaques

**DOI:** 10.1101/2023.08.30.555589

**Authors:** Sam K. Patterson, Ella Andonov, Alyssa M. Arre, Melween I. Martínez, Josué E. Negron-Del Valle, Rachel M. Petersen, Daniel Phillips, Ahaylee Rahman, Angelina Ruiz-Lambides, Isabella Villanueva, Amanda J. Lea, Noah Snyder-Mackler, Lauren J.N. Brent, James P. Higham

**Author notes:** Corresponding author: Sam Patterson, 25 Waverly Pl, New York, New York 10003 USA.

## Abstract

Exposure to adversity during early life is linked to lasting detrimental effects on evolutionary fitness across many taxa. However, due to the challenges of collecting longitudinal data, especially in species where one sex disperses, direct evidence from long-lived species remains relatively scarce. Here we test the effects of early life adversity on male and female longevity in a free-ranging population of rhesus macaques (*Macaca mulatta*) at Cayo Santiago, Puerto Rico. We leveraged six decades of data to quantify the relative importance of ten forms of early life adversity for 6,599 macaques (3,230 male, 3,369 female), with a smaller sample size (N=299) for one form of adversity (maternal social isolation) which required high-resolution behavioral data. We found that individuals who experienced more early life adversity died earlier than those who experienced less adversity. Mortality risk was highest during early life, defined as birth to four years old, suggesting acute survival effects of adversity, but heightened mortality risk was also present in macaques who survived to adulthood. Females and males were affected differently by some forms of adversity, and these differences might be driven by varying energetic demands, female philopatry, and male dispersal. By leveraging data on thousands of macaques collected over decades, our results show that the fitness consequences of early life adversity are not uniform across individuals but vary as a function of the type of adversity, timing, and social context, and thus contribute to our limited but growing understanding of the evolution of early life sensitivities in long-lived species.

Significance Statement

Exposure to early life adversity, even when conditions subsequently improve, can have profound and persistent consequences for human health. Negative effects of early life adversity appear widespread across the animal kingdom. To date, however, direct evidence from long-lived species is relatively scarce due to the difficulties of collecting data from early life till death. We leverage six decades of observations on thousands of free-ranging male and female rhesus macaques to examine the complex ways early life adversity impacts survival. Our results suggest that the type of adversity and life history factors intersect to impact immediate and downstream survival. By studying early life adversity across environments, cultures, contexts, and species, we can better understand the evolutionary underpinnings of early life sensitivities.

## Introduction

Exposure to adversity such as food shortages and social isolation during early life can result in long-term health and evolutionary fitness consequences in a wide range of species such as insects, birds, fish, reptiles, and mammals (Cooper & Kruuk, 2018; Eyck et al., 2019; Lea & Rosenbaum, 2020; Lu et al., 2019). For example, female red deer (*Cervus elaphus*) that face high resource competition during development exhibit accelerated senescence in adulthood (Nussey et al., 2007). Early life adversity in female baboons (*Papio sp.*) is associated with reduced fecundity and poorer offspring survival (Lange et al., 2023; Lea et al., 2015; Patterson et al., 2021; Tung et al., 2016, 2023; Weibel et al., 2020; Zipple et al., 2019). Organisms are hypothesized to adjust their developmental trajectories in response to early life adversity in order to improve immediate survival (Lea et al., 2017; Lea & Rosenbaum, 2020; Patterson, Petersen et al., 2023), but such adjustments may lead to these detrimental outcomes in adulthood. A small but growing number of studies have tested the long-term impacts of early life adversity, but further research on how different forms of early life adversity shape the timing of fitness consequences in a variety of species, populations, and contexts is needed to better understand the evolution of early life sensitivities to adversity.

Survival, which is a prerequisite for reproductive success and an important aspect of evolutionary fitness, has been linked to early life adversity in a number of species. Given the difficulty of measuring fitness directly, a common approach is to quantify lifetime reproductive success (LRS), which is the total number of offspring produced over an individual’s lifetime (Clutton-Brock 1988). Adult lifespan is the biggest contributor to LRS in long-lived species like roe deer (*Capreolus capreolus*), baboons, rhesus macaques (*Macaca mulatta*) (Blomquist, 2009), and gorillas (*Gorilla beringei beringei*) (Clutton-Brock et al., 1983; Kjellander et al., 2004; Rhine et al., 2000; Robbins et al., 2011; Van de Walle et al., 2022). Females who live longer have a longer reproductive span, and are able to produce more offspring. Long-term studies have shown that early life adversity is associated with reduced survival in adult female baboons and hyenas (*Crocuta crocuta*) (Gicquel et al., 2022; Lange et al., 2023; Strauss et al., 2020; Tung et al., 2016). In populations characterized by high mortality rates prior to reproductive maturity, LRS based on the total number of offspring *reaching reproductive maturity* is the best proxy for fitness (e.g., Alif et al. 2022). Death prior to maturity has severe fitness costs for the organism who fails to reproduce, but also for the organism’s parents. Male and female gorillas exposed to early life adversity experience reduced survival prior to reproductive maturity, but do not experience survival costs after maturity (Morrison et al., 2023). As such, adult survival patterns in gorillas differ from those in yellow baboons and spotted hyenas, but pre-reproductive survival patterns are not yet available across species for comparison. More studies across different species are thus needed to both identify the fitness consequences of early life sensitives and to draw comparisons and better understand the evolutionary pressures which shape the developmental responses to adversity and ultimately influence fitness across the lifespan.

The fitness consequences of early life adversity might vary in a sex-dependent manner due to differences in life history strategies. During adverse early life conditions, the sex with more energetically demanding traits is predicted to be more susceptible to nutritional constraints (Clutton-Brock, 1994; Clutton-Brock et al., 1985). Further, during adverse conditions, parents are predicted to reduce investment in more energetically costly offspring, thus exacerbating consequences of adversity especially during the period of care (Clutton-Brock et al., 1985; Trivers & Willard, 1973). In many species, males are considered more energetically costly given faster growth and larger body size compared to females. When male fitness is largely determined by access to mates via competitive ability, males should also invest in costly developmental processes like play and motor skill development (Lonsdorf, 2017). In support of these predicted differences, among red deer (*Cervus elaphus*), maternal death prior to weaning was linked to higher mortality risk among male compared to female offspring (Andres et al., 2013). Sex-dependent effects of early life adversity are challenging to study because many species are characterized by sex-biased dispersal such that pre– or post-dispersal data are typically missing for individuals of the dispersing sex. More studies are thus needed that can follow both males and females from birth till death to investigate how life history and parental investment strategies shape fitness consequences and developmental responses to early life adversity.

Here we leverage a large historical dataset of the free-ranging rhesus macaques (*Macaca mulatta*) of Cayo Santiago to advance our understanding about the magnitude, form, timing, and sex-dependence of early life adversity effects. Complete demographic records extend back to the 1960s for thousands of male and female macaques at the Cayo Santiago field site. In this population, abusive maternal care behavior is linked to differences in HPA function in juveniles, the presence of a competing younger sibling is linked to reduced survival during juvenility, and exposure to hurricanes and high population density during early life are linked to life history trade-offs in adulthood (Lee et al., 2019; Luevano et al., 2022; Petrullo et al., 2016). We examine the effects of ten forms of potential early life adversity on sex-specific mortality risk across early life, and separately, across adulthood. By examining mortality across the lifespan, we can identify the full extent of variation in the fitness consequences of early life adversity. If mortality consequences are severe in early life, this might influence survival patterns observed in adulthood (e.g., survivorship bias), and it would be important to consider why some individuals can survive to reproductive debut while others do not.

We predict that rhesus macaques exposed to greater amounts of early life adversity will have increased mortality risk. We predict mortality risks will be more severe during the first four years of life when individuals are still growing and adversity is more recent. However, we also predict heightened mortality risk will persist into adulthood among those who survive past four years of age. Given male life history strategies in this species (Hoffman et al., 2008) that prioritize costly traits like faster growth, larger body size, and motor skill development (Kulik et al., 2015; Schwartz & Kemnitz, 1992; Turcotte et al., 2022), we predict that early life adversity will exert larger effects on males than females.

## Methods

### Study site and population

We studied a free-ranging population of rhesus macaques living on Cayo Santiago, a 15.2-ha island off the southeastern coast of Puerto Rico. The current population of individually recognized ∼1,700 rhesus macaques are the descendants of 409 monkeys that were transported from India to the island in 1938. This population is managed by the Caribbean Primate Research Center (CPRC) of the University of Puerto Rico. Monkeys are fed monkey chow daily and water catchments provide *ad libitum* access to drinking water. The monkeys live in naturally forming multi-male, multi-female social groups characterized by dominance hierarchies and male dispersal. Monkeys mate with multiple partners and breed seasonally. This species is characterized by sexual dimorphism with males exhibiting larger body mass and canine length than females (Schwartz & Kemnitz, 1992; Turcotte et al., 2022). Males queue for dominance rank, have large testes, and experience indirect male-male competition (Higham & Maestripieri, 2014; Kimock et al., 2019, 2022). Male mortality is highest during the mating season, consistent with the notion that males prioritize investment in mating effort (Higham & Maestripieri, 2014; Hoffman et al., 2008). Females prioritize investment in gestation and lactation, and face the highest mortality risk during the birth season (Hoffman et al., 2008). The island is free of predators and there is no regular veterinary intervention, so the primary causes of death are illness and injury (Pavez-Fox et al., 2022).

During the study period (1960-2021), observers monitored and recorded demographic events daily. These records include births, deaths, sex, matriline, matriline rank, maternal identification, sires when genetic data were available, and group emigration and immigration events. A genetic pedigree is available for much of the population (Widdig et al., 2016). Daily total rainfall and mean maximum temperature data were pulled from the NOAA station in Rio Piedras, Puerto Rico. Over a 61-year period (1959-2020), data were not recorded by this NOAA station for 21% of days. Rather than removing a large portion of data, we imputed missing rainfall and temperature data using the ‘mice’ package in R (Buuren & Groothuis-Oudshoorn, 2011). This study includes 6,599 individuals for which there are complete data available. We had data covering the entire lifespan–birth to death–from 2,513 macaques. The remaining 4,086 macaques were either alive at the time of this study or were removed from the island prior to natural death as a result of population management (i.e., right-censored samples).

### Early life adversities

We used historical demographic records to assess individual exposure to early life adversity (Tung et al., 2016). We consider ten forms of potential early life adversity based on previous research on this population and other species, as described below. In choosing time periods of exposure for each form of adversity, we followed Tung and colleagues 2016:

### Maternal loss

Maternal death increases offspring mortality in humans and other mammals (Cayo Santiago macaques: Blomquist, 2013; red deer: Andres et al., 2013; Asian elephants (*Elephas maximus*): Lahdenperä et al., 2016; humans: Sear & Mace, 2008; chimpanzees (*Pan troglodytes*): Stanton et al., 2020; yellow baboons: Tung et al., 2016). Following previous work (Tung et al., 2016; Zipple et al., 2021), we consider an individual to experience maternal loss if their mother died (including natural death (N=1,165) and permanent removal from the population (N=299)) before the individual reached 4 years of age. This four year period includes the period during which young macaques are nutritionally and socially dependent on their mothers. While a mother’s removal from the population had to occur while the offspring was alive (i.e., prior to offspring death if they have died) to be considered an adversity, this was not a requirement for maternal death because an imminent maternal death is linked to offspring mortality risk–an association likely explained by poor maternal condition (Zipple et al., 2021). If maternal loss occurred after the offspring reached four years old, we did not consider the offspring to have experienced this source of early life adversity. Maternal loss was measured as a binary variable: experienced maternal loss or did not experience loss.

### Competing sibling

The presence of a close in age younger sibling represents a source of competition over maternal resources and is associated with higher mortality risk (Cayo Santiago macaques: Lee et al., 2019; yellow baboons: Tung et al., 2016). We considered a sibling to be a competitor if the sibling was born within 355 days, which represented the bottom quartile of interbirth intervals in our sample. Last born offspring and individuals which died before their sibling was born did not experience this adversity. The presence of competing siblings was measured as a binary variable.

### Group size

High group size and high population density are indicative of more competition and are associated with reductions in fecundity (Cayo Santiago macaques: Luevano et al., 2022; red deer: Clutton-Brock et al., 1982, 1983; meerkats (*Suricata suricatta*): Clutton-Bock 2008). We used group size as a proxy for within-group competition. Demographic records were used to construct group composition over the study period. Group size was defined as the number of adults (>=4 years of age) of both sexes in an individual’s social group on the day that individual was born (Tung et al., 2016), and was included in our models as a continuous variable (range: 2-222 individuals).

### Primiparity

The high energetic demands on first time mothers can result in negative outcomes for offspring such as increased mortality risk (Rhesus macaques: Bercovitch et al., 1998; Blomquist, 2013;Nuñez et al., 2015; vervet monkeys (*Chlorocebus sabaeus*): Fairbanks & Mcguire, 1995; mantled howler monkeys (*Alouatta palliata*): Glander, 1980; Asian elephants: Mar et al., 2012; olive baboons (*Papio anubis*): Smuts & Nicolson, 1989). We used a binary measurement for primiparity: first born or not first born.

### Matriline rank

Dominance rank mediates access to food and is linked to survival, fecundity, and offspring growth (Cayo Santiago macaques: Blomquist et al., 2011; Weiß et al., 2016; yellow baboons: Altmann & Alberts, 2005; olive baboons: Garcia et al., 2009; chacma baboons (*Papio ursinus*): Johnson, 2003). Matrilineal dominance hierarchies for a given social group and year are recorded as categorical – high, middle, low – based on data from Donald S. Sade and John D. Berard, who recorded dyadic agonistic interactions (e.g., threats, displacements, submissive behaviors) across the year to calculate matrilineal dominance matrices (Lee et al., 2019; Missakian, 1972).

### Kin network

Among prime aged adult females at Cayo Santiago, the presence of more maternal kin is linked to better survival in any given year (Brent et al., 2017). We measured an individual’s maternal kin network size at birth as the number of living females over 4 years of age with a relatedness coefficient of at least 0.063. This relatedness coefficient was chosen because 0.063 represents the threshold at which macaques in this population can recognize kin via vocalizations (Rendall et al., 1996), and this threshold was used in previous work showing a positive association between the number of relatives present and adult survival (Brent et al., 2017). Kin network size was included as a continuous variable (range: 1-21 individuals).

### Maternal social connectedness

Greater social connectedness is associated with better survival and better offspring survival (Cayo Santiago macaques: Brent, Heilbronner, et al., 2013; Ellis et al., 2019; yellow & chacma baboons: Archie et al., 2014; Silk et al., 2003, 2010). We used behavioral data collected during 10-min focal animal samples on adults in several social groups from 2010-2017 (details provided in the SOM). To measure maternal social connectedness, we calculated a composite sociality index (CSI) using the affiliative social behaviors, approaches and grooming. For each mother in each year, we tabulated the rate of approaches (approaches to and from other adult females / hours observed) and the rate of grooming bouts (number of grooming bouts given and received / hours observed). A mother’s approach and grooming rates were divided by the mean rate for all adult females in each social group in each year. These standardized approach and grooming rates were added together and divided by 2 (the number of behaviors) to create the CSI for each mother. Here, we followed Tung and colleagues (2016): for each offspring in our analyses, we averaged their mother’s composite sociality index for the first two years of life.

### Rainfall

More rainfall is indicative of greater food and water availability, and is linked to greater fecundity and better survival (primates: Campos et al., 2017; yellow baboons: Lea et al., 2015; gelada monkeys (*Theropithecus gelada*): Sloan et al., 2022). However, because food and water are provisioned and due to the negative effects of tropical storms at Cayo Santiago, low rainfall might not be as relevant or have negative consequences in this population. Here, we used total rainfall across the first year of life (range: 1,021.4-3,157.1 mm) (Tung et al., 2016).

### Temperature

Higher temperatures are linked to fecundity, cognition, and mortality (Western Australian magpies (*Cracticus tibicen dorsalis*): Blackburn et al., 2022; southern pied babbler (*Turdoides bicolor*): Bourne et al., 2020; dairy cattle (*Bos taurus*): Polsky & von Keyserlingk, 2017; gelada monkeys: Sloan et al., 2022). Here, we averaged mean maximum daily temperatures across the first year of life (range: 85.12-89.89 F) (Tung et al., 2016).

### Hurricanes

Exposure to major hurricanes is linked to female reproductive strategies, demographic roles, and immunological aging (Cayo Santiago macaques: Diaz et al., 2023; Luevano et al., 2022; Watowich et al., 2022). We recorded individual exposure to any of the 3 major hurricanes that had major impacts on Cayo Santiago (Hugo on September 18, 1989, Georges on September 21, 1998, and Maria on September 20, 2017) during the first year of life. Hurricane exposure was not included in previous studies of early life adversity, so we chose the first year of life as our window of exposure to align with our other weather variables, rainfall and temperature. If individuals were exposed to hurricanes when they were over one year old, we did not consider them to have experienced this adversity as *early life* adversity.

Several approaches have been used to conceptualize, process, and analyze early life adversity data. Some studies of early life adversity use broad, cumulative measures of adversity, while others focus on different forms of adversity separately (Gunnar, 2020; Smith & Pollak, 2020). Empirical evidence suggests that the accumulation of multiple adversities is a better predictor of adult outcomes than any particular form of adversity, but there is also evidence that specific forms of adversity lead to different outcomes (Gicquel et al., 2022; Gunnar, 2020; Tung et al., 2016). Here, we examine cumulative adversity measures and then examine individual forms of adversity separately. To construct a cumulative early life adversity index, we summed individuals’ exposure to different forms of adversity. Previous studies typically relied on binary scores for each form of adversity. Here, we use continuous measures of adversity when feasible. For purposes of the cumulative index, continuous measures (i.e., group size, kin network size, high temperature, and rainfall) were normalized so values range from zero to one. For binary measures (i.e., maternal loss, being a first born, presence of a competing sibling, and hurricane exposure), individuals were assigned a value of one if they experienced a given form of adversity and a value of zero if they did not experience that form of adversity. Those born into high ranking matrilines were assigned a zero, mid ranking matrilines were assigned 0.5, and low ranking matrilines were assigned a value of one. As such, each variable ranged from zero to one and were summed together into a cumulative index to represent the total exposure to early life adversity. Our main cumulative early life adversity index could range from 0-9 because it included nine variables: maternal loss, presence of a competing younger sibling, high group size, primiparity, low matrilineal dominance rank, small kin network, hurricane exposure, high temperature, and low rainfall. Maternal social connectedness is not included because it was derived only for a subset of our data.

## Data analysis

To determine if early life adversity predicts survival, we used survival models. The outcome variable was age at death. Individuals who were either still alive at the end of the study or removed from the island for population control were right-censored. We ran models on the full sample of all ages (N=6,599) but right-censored to four years old to examine early life mortality, and we ran models on a subsample of individuals who survived beyond four years of age (N=2,866) to examine mortality across adulthood. Early life adversity predictor variables were modeled two ways: 1) cumulative index models include all forms of adversity summed together into one variable, and 2) multivariate models include each form of adversity modeled as individual predictor variables. We ran separate models to examine the survival effects of maternal social connectedness during early life because our focal behavioral data does not span the entire study period (N=299 early life survival; N=101 adult survival). Models included early life adversity index, sex, and an interaction term between sex and early life adversity. Models also included a varying intercept for birth year and maternal identification. The different forms of adversity we examined were not correlated, but high temperatures in the first year of life were highly correlated with birth year (Table S1).

We first used Cox survival models, but the proportional hazards assumption in the Cox model was violated in three out of four models. Specifically, early life and adult multivariate models and the early life cumulative index model violated assumptions. Individual sex, maternal loss, high temperatures, group size, kin network size, matrilineal rank, hurricane exposure, and four of the sex interaction terms violated proportional hazard assumptions (cox.zph function in R package, “survival”: p<0.05). Instead, we fit Accelerated Failure Time (AFT) survival models with a Weibull distribution. The presence of a competing younger sibling is time-dependent since individuals cannot experience this exposure unless they survive till a given age, i.e., until it’s biologically possible for the mother to give birth again. To include this variable, we would need to include it as a time-varying variable in a Cox proportional hazard model. As such, we were unable to test how the presence of a competing younger sibling affects survival in early life and we excluded this variable from the cumulative index for the early life survival model. We could, however, examine this in adulthood since all individuals in the sample survived to adulthood and the presence of a competing younger sibling is not time-varying.

Genetics can contribute to the effects of early life adversity. For example, individuals experiencing maternal loss might have shorter lifespans due to genes shared by both the mother and offspring. To estimate to what extent variance in survival is explained by genetics and early life adversity, we account for pedigree in a subsample of the data for which we had complete pedigree information (N=923 individuals during early life; N=307 adults). To do so, we used the animal model and incorporated the relationship covariance matrix as a random effect (Wilson et al., 2010). The models with and without pedigree produced similar results (Table S2). The effects of pedigree on survival were substantial, but accounting for genetic relatedness in the model did not diminish the effects of early life adversity on survival in early life (with pedigree: ꞵ=-0.33±0.12; without pedigree: ꞵ=-0.31±0.11) or adulthood (with pedigree: ꞵ=-0.12±0.04; without pedigree: ꞵ=-0.12±0.04). The full pedigreed sample is smaller than our main dataset because paternity is unknown for many animals earlier in the study. Because the effects of early life adversity were unaffected by pedigree inclusion and because the sample size for pedigree inclusion is much smaller, we have presented the larger set of data without pedigree in the main text.

Models were run with the brms package (v 2.16.3) in R (v 4.1.2) (Bürkner, 2017). All continuous predictor variables were transformed to a mean of 0 and a standard deviation of 1. All models are Bayesian, and we used weakly informative priors for fixed effects, setting the mean to zero and the standard deviation to one. To produce more accurate predictions for age at death, we used more regularizing priors for the intercept (a mean of 1 and standard deviation of 0.1 for the early life survival models, and a mean of 12 and standard deviation of 0.4 for the adult survival models). Specifically, our analyses contain a high proportion of right-censored cases, which can lead to model predictions that overestimate life expectancy (Alam et al., 2022). We use credible intervals to determine whether the effect of a variable is substantial or not. If the 85% credible interval for an effect does not overlap with zero, the effect is considered substantial. When the vast majority of the 85% credible interval does not span zero, but there is some overlap, we consider the model to be uncertain about the effect. The code and data used can be found here: https://github.com/skpatter/ELA_Survival_Macaques

## Results

### Cumulative early life adversity is associated with reduced survival during early life and during adulthood

Individuals who experienced more cumulative early life adversity had higher mortality during early life (ꞵ=-0.29±0.07; Figure 1, Table S3). There were no clear differences in mortality for males versus females during early life (ꞵ=-0.05±0.10), and there was no evidence that early life adversity differentially affected mortality risk as a function of sex during early life (ꞵ=-0.07±0.09; Figure 1, Table S3). Adults who experienced more cumulative early life adversity had shorter lives than adults with less early life adversity (ꞵ=-0.04±0.02; Figure 1, Table S3). Among adults, females lived longer than males (ꞵ=-0.13±0.02), and there was no evidence that cumulative early life adversity differentially affected mortality risk among males and females (ꞵ=0.00±0.02; Figure 1; Table S3).

**Fig 1.**
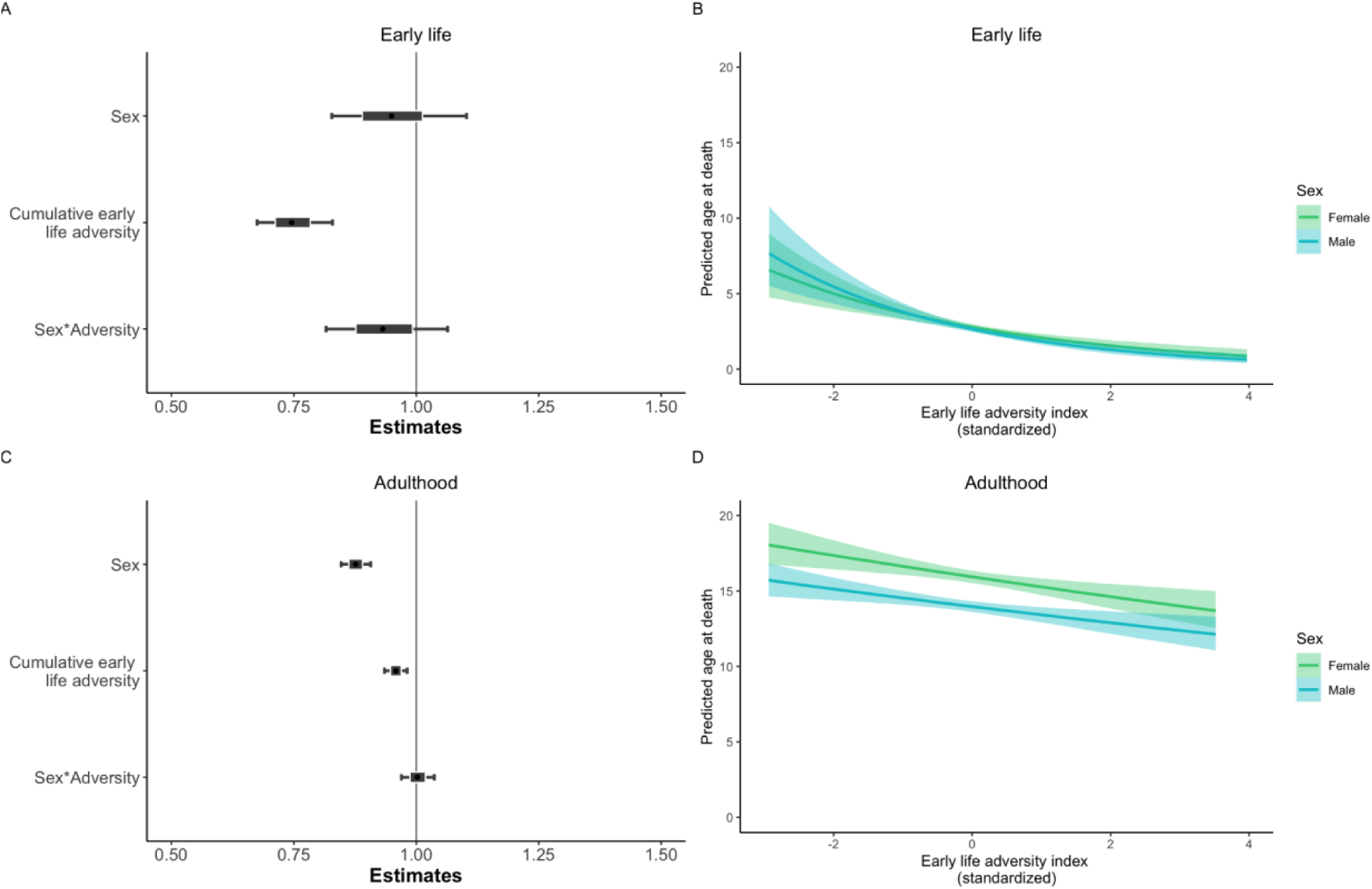
Model effects of cumulative early life adversity and sex on survival during early life (A) and adulthood (C). The outer bars show the 85% credible intervals, the inner box shows the 50% credible intervals, and the black circle in the middle shows the median of the posterior distribution. Model predictions are shown for the effect of cumulative early life adversity on lifespan in early life (B) and adulthood (D). Green predictions represent females and blue predictions represent males. The solid lines show the median estimates and the shaded region shows the 85% credible intervals.

Between those who experienced the least and the most amount of cumulative early life adversity in our sample, these effects translate to a 4.78-year difference in average life expectancy among adult females and a 3.94-year difference in average life expectancy among adult males.

### Several forms of early life adversity are associated with reduced survival, and some effects are sex-dependent

Individuals who **lost their mother** during the first four years of life had a higher mortality risk during early life (ꞵ=-0.33±0.14) and adulthood (ꞵ=-0.06±0.04) than those who did not lose their mother (Figure 2, Table S4). Maternal loss had a larger negative effect on sons than daughters in both early life (ꞵ=-0.32±0.20) and adulthood (ꞵ=-0.05±0.06; Figure 3, Table S4). **Higher temperatures** during the first year of life were associated with higher mortality rises in early life (ꞵ=-0.06±0.24) and adulthood (ꞵ=-0.02±0.02), but the models were uncertain about these effects (Figure 2, Table S4). The effect of high temperatures on survival was moderated by sex (Figure 3; Table S4). Higher temperatures during the first year of life were more strongly associated with reduced survival among males than females during early life (ꞵ=-0.23±0.09), but in adulthood, only females experienced this survival cost (ꞵ=0.02±0.03).

**Fig 2.**
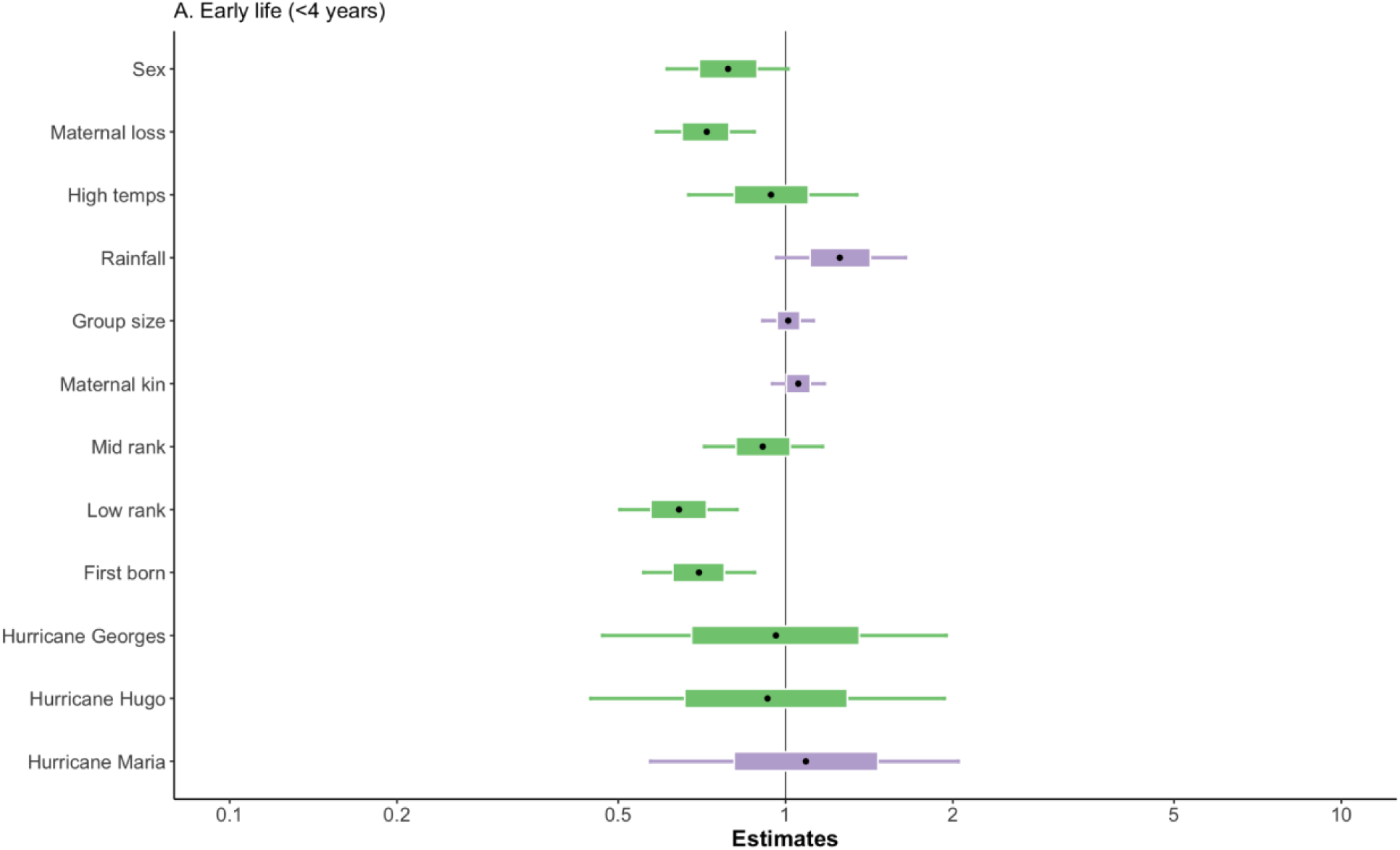
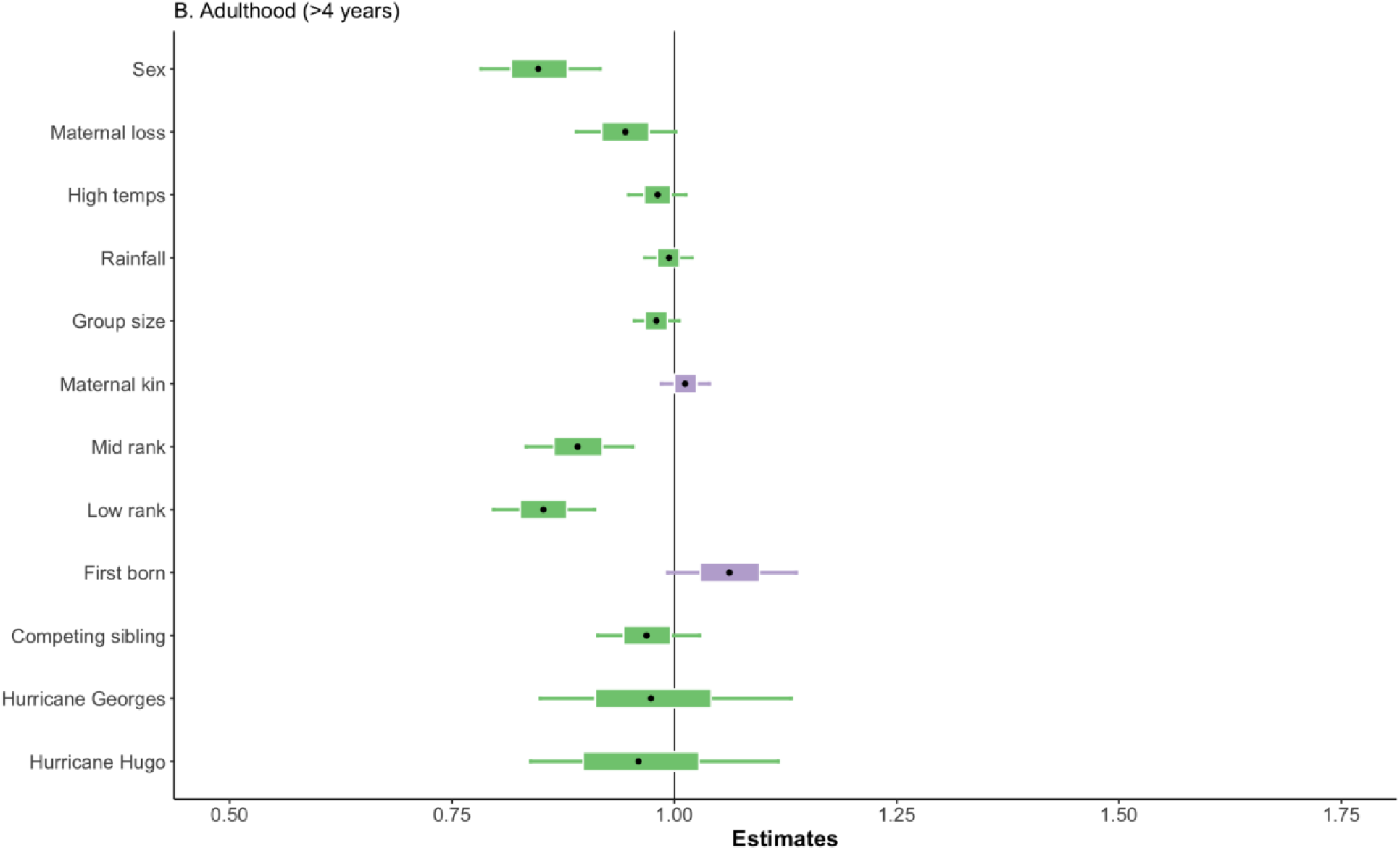
Model effects of sex and the forms of early life adversity on survival during early life (A) and adulthood (B). The outer bars show the 85% credible intervals, the inner boxes show the 50% credible intervals, and the black circles in the middle show the medians of the posterior distributions. Green shading represents negative effect sizes, meaning that the variable is associated with shorter lifespans, and purple shading represents positive effect sizes, meaning that the variable is associated with longer lifespans. For interaction effects, see Figure S1.

**Fig 3.**
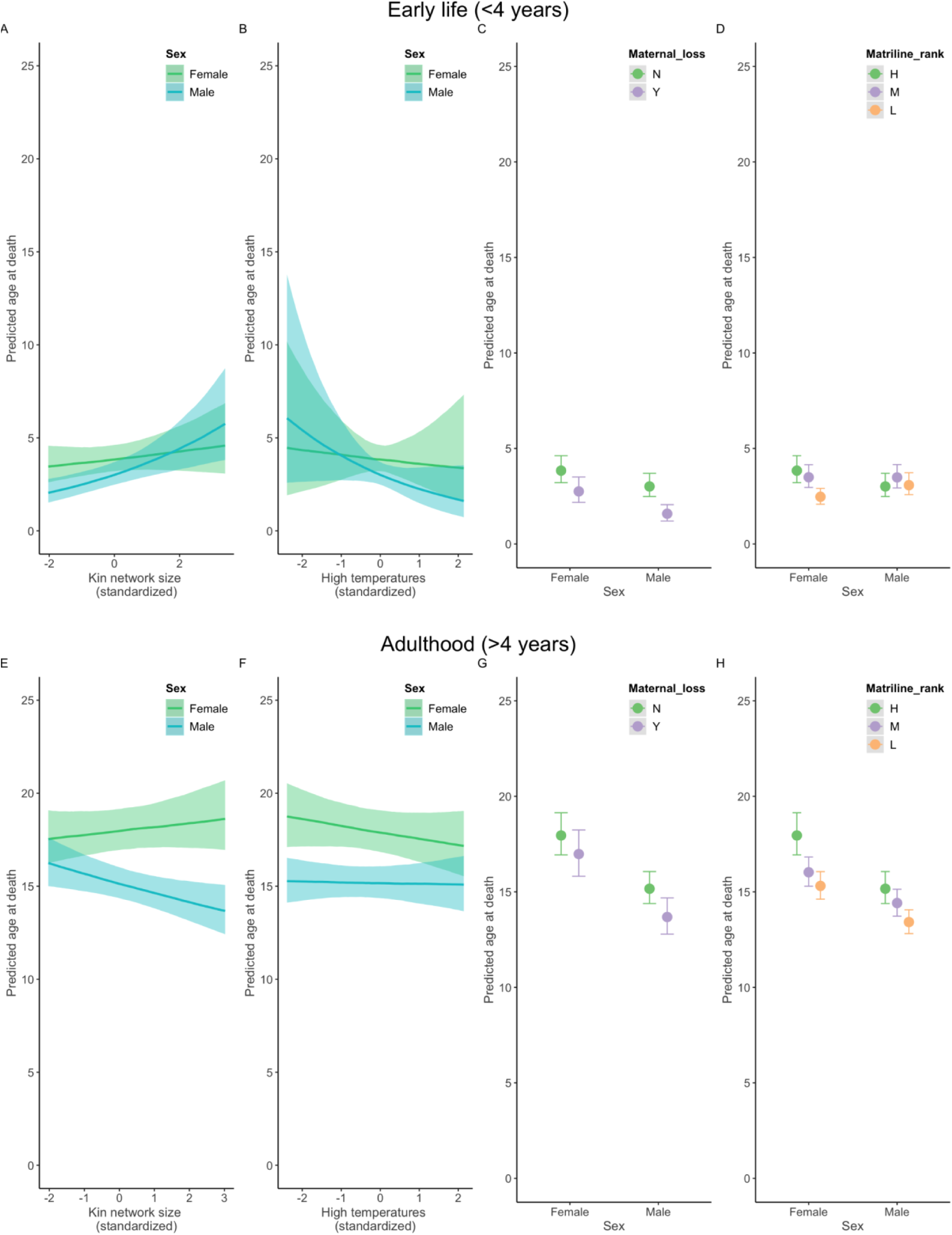
Interactions between sex and three forms of early life adversity on adult survival. (A) and (E) Predicted relationship between maternal kin network size at birth and survival for females (green) and males (blue). (B) and (F) Predicted relationship between high temperatures during the first year of life and survival for females (green) and males (blue). (C) and (G) Predicted relationship between maternal loss during the first four years of life and survival for males versus females. The circles show the median estimate, and the bars show the 85% credible intervals. (D) and (H) Predicted relationship between matrilineal rank and survival for males versus females. The circles show the median estimate, and the bars show the 85% credible intervals.

**First-born** offspring had elevated mortality risk during early life (ꞵ=-0.36±0.16). In contrast, individuals born to primiparous mothers had better survival in adulthood than those born to multiparous mothers, although the model was uncertain about this effect (ꞵ=0.06±0.05). Effects of maternal primiparity were not moderated by sex (early life: ꞵ=0.09±0.22; adulthood: ꞵ=-0.04±0.07). Macaques born into **low ranking matrilines** had a higher mortality risk during early life (ꞵ=-0.44±0.17) and adulthood (ꞵ=-0.16±0.05) than those born into high ranking matrilines. Matrilineal rank was more strongly associated with female survival than male survival during both periods of life (early life: ꞵ=0.47±0.22; adulthood: ꞵ=0.04±0.06; Figure 3; Table S4). We also treated matrilineal rank as an ordinal variable and found similar results (Table S4).

**Smaller maternal kin networks** at birth were associated with higher early life mortality risk, especially for males (ꞵ=0.05±0.07; sex interaction: ꞵ=0.14±0.10). Smaller maternal kin networks at birth were associated with better survival for adult males, but reduced survival for adult females (ꞵ=0.01±0.02; sex interaction: ꞵ=-0.05±0.03; Figure 3; Table S4). The presence of **competing younger siblings** was associated with higher mortality risk, but the model was uncertain about this effect (ꞵ=-0.03±0.04). Although the model was uncertain, a competing sibling had a slightly larger effect on females (ꞵ=0.06±0.06). We were unable to examine survival effects during early life given time-varying issues.

Low **rainfall** was associated with reduced survival during early life, but the model was uncertain about this estimate (ꞵ=0.23±0.18). No effect of rainfall was found during adulthood (ꞵ=-0.01±0.02; Figure 2; Table S4). No effect of **group size** was observed during early life (ꞵ=0.01±0.07), and while the model was uncertain, it seems adults born into larger groups exhibited reduced survival (ꞵ=-0.02±0.02; Figure 2; Table S4). No effect of major **hurricanes** was observed during early life (Georges: ꞵ=-0.04±0.50; Hugo: ꞵ=-0.08±0.50; Maria: ꞵ=0.08±0.44) or adulthood (Georges: ꞵ=-0.02±0.10; Hugo: ꞵ=-0.04±0.10; Figure 2; Table S4). The model was uncertain about the effects of maternal social isolation; individuals born to **socially isolated mothers** seemed to have higher mortality during early life than those born to more socially connected mothers (ꞵ=0.15±0.17; Table S5), and males were more affected by this than females (ꞵ=0.29±0.25). No effect of maternal social isolation was observed among adults (ꞵ=-0.04±0.02; Table S5). Model comparisons revealed no substantial difference between models constructed with the cumulative early life adversity index versus those constructed with each form of early life adversity separately (Table S6).

## Discussion

Our findings indicate that early life adversity shapes both early life survival and adult survival in free-ranging rhesus macaques. Individuals experiencing more cumulative early life adversity lived shorter lives than those with less adversity. The effect size of early life adversity on mortality risk was larger in the first four years of life than adulthood, but risks were also elevated in adulthood. Strong effects on early life mortality risk are consistent with the notion of an overall greater vulnerability during development (Walasek et al., 2022; West-Eberhard, 2003). Given the fitness costs of dying prior to reproduction, our results demonstrate that the effects of early life adversity prior to maturity have major fitness ramifications and the full consequences of early life adversity are likely to be larger than predicted in previous studies focused on adult fitness.

The various forms of potential adversity we measured did not contribute equally to survival with maternal-related adversities exhibiting the largest effects. Maternal death in the first four years of an individual’s life and low matrilineal rank were associated with higher mortality risk in early life and adulthood. The lasting effects of these maternal-related adversities are unsurprising given similar consequences in other mammalian species (e.g., Stanton et al., 2020; Strauss et al., 2020; Tung et al., 2016; Zipple et al., 2019), as well as consequences of parental-related hardships in humans (Fields et al., 2021; Glover et al., 2018; O’Donnell et al., 2014; Reid et al., 2018; Thayer & Kuzawa, 2014). Other maternal effects were also linked to mortality, but the effects were not as strong or straightforward. Survival advantages were observed among offspring born to more socially connected mothers, but there was considerable variation in this effect and it did not persist into adulthood. Future work is needed to examine how an individual’s own social connectedness in adulthood potentially interacts with the maternal social network they experienced during early life to shape survival. Small maternal kin networks, a proxy for low social support, predicted reduced survival, but effects were sex-dependent. From previous analyses, we know the presence of a competing younger sibling increases mortality risk during early life (Lee et al., 2019), and these effects appear to persist into adulthood, at least for females. Offspring born to first time mothers were more likely to die during early life than those who were not the first born. However, while our model estimates were uncertain, among those who survived into adulthood, survival odds were likely better for those born to primiparous than multiparous mothers. Given the strong negative effects of primiparity on offspring survival in the first four years of life, the higher survival odds of adults who were first borns could reflect survivorship bias. Alternatively, confounding variables that were not included in the analyses such as the presence of the grandmother, mother, and other kin across adulthood might also contribute to the patterns observed among adults.

Other forms of adversity, which were not directly related to the maternal environment, also contributed to survival. High temperatures generally predicted reduced survival, but this effect was sex-dependent. High temperatures are not typically included in measures of adversity for primates, but identifying potential health consequences and the underlying mechanisms is of interest given rising global temperatures (Hondula et al., 2015). While low rainfall seemed to have some negative effects on survival during early life, we found no substantial relationship between rainfall and adult survival. High group size at birth was not associated with survival during early life, but seemed to have some negative effects on adult survival. Rainfall and group size might have limited effects on survival because drinking water and food are provisioned in the study population. While the macaques still compete over access to food and water resources, competition is likely reduced compared to wild populations.

Consistent with previous analyses in this population (Luevano et al., 2022; Morcillo et al., 2020), we did not find substantial impacts of early life hurricane exposure on survival. This is surprising given that macaques in this population exposed to Hurricane Maria showed divergent immune cell gene regulation, suggestive of accelerated aging (Watowich et al., 2022). Exposure to major hurricanes also led to greater heterogeneity in reproductive strategies and longevity, and macaques might reduce fertility as a strategy to prioritize survival odds (Diaz et al., 2023; Morcillo et al., 2020). Given our results showing heat effects on mortality and the fact that temperatures increase following hurricanes on Cayo Santiago (Testard et al., 2021; Watowich et al., 2022), hurricanes might affect macaques indirectly via factors such as deforestation, shade scarcity, and heat. Further, given the recency of Hurricane Maria and our small sample of individuals exposed to Maria in this study’s dataset, we are currently limited in our ability to analyze survival outcomes for this most recent hurricane event. Potential impacts of Hurricane Maria may also have been socially buffered – macaques on Cayo Santiago adjusted their social networks after Hurricane Maria (Testard et al., 2021) and built new social connections, which may buffer negative impacts.

The survival effects of some forms of early life adversity were sex-dependent. Studies of early life adversity in long-lived animals have typically only been assessed in one sex, the non-dispersing sex (e.g., Gicquel et al., 2022; Patterson et al., 2022; Tung et al., 2016), but studies that have examined males and females produced mixed results. While male red deer are more negatively impacted by maternal death than female deer (Andres et al., 2013), there do not appear to be any sex-based differences in susceptibility to the survival costs of cumulative early life adversity in gorillas (Morrison et al., 2023), though males and females might vary in their responses to specific types of adversity. We predicted that early life adversity would have greater effects on mortality risk in males than females because of sex differences in life history strategies with males typically prioritizing more energetically intensive processes (Higham & Maestripieri, 2014; Hoffman et al., 2008; Schwartz & Kemnitz, 1992; Turcotte et al., 2022). We found some evidence in support of this prediction. During early life, male survival was more negatively affected by three forms of adversity: small maternal kin networks, high temperatures, and maternal loss. In adulthood, males continued to suffer greater costs of early maternal loss, perhaps reflecting the long-term costs of severe energetic constraints during early life. Males might be more affected by these adversities than females prior to reproductive maturity due to their energetically costly developmental trajectories and/or due to maternal decisions to reduce investment in energetically costly offspring during harsh environments (Clutton-Brock, 1994; Clutton-Brock et al., 1985; Trivers & Willard, 1973). Further research is needed to investigate how effects of early life adversity might be moderated or mediated by developmental trajectories and parental investment strategies.

We also found evidence contrary to our predictions about sex-dependent effects: in adulthood, females were more affected by several forms of early life adversity than males. These findings are consistent with studies on later life mortality in humans which suggest that women tend to be more susceptible to early life adversity than men (Johnson et al., 2020; Lee & Ryff, 2019). Differences between men and women could be due to survivorship bias, underreporting of symptoms among men, confounds of societal gender biases and inequities, or due to underlying biology (Johnson et al., 2020). Given the complexity of potential factors contributing to effects in humans, it can be useful to turn to simpler animal models. Experimental work on mice also showed that early life adversity resulted in increased anxiety and depressive behaviors in female mice but not males (Bondar et al., 2018; Goodwill et al., 2019). Our results suggest species-specific social patterns and underlying biology might contribute to sex-dependent patterns associated with early life adversity. First, adult females were more affected by matriline rank than adult males likely because males disperse (Weiß et al., 2016), female dominance hierarchies are fairly stable across time (Blomquist et al., 2011), and females typically inherit dominance rank via their matriline. That this sex difference was already apparent in early life might reflect variation in social networks and social priorities between immature males and females (Amici et al., 2019). Second, being born into large maternal kin networks had a positive effect on adult female survival but a negative effect on adult male survival. Given dispersal, males might not receive any immediate benefits of kin support in adulthood and thus only experience the long-term costs associated with earlier competition, consistent with the idea that individuals face tradeoffs between benefits of kin support and costs of competition with kin (Croft et al., 2017). Third, males were more susceptible than females to high temperatures during early life, but females were more susceptible in adulthood. Sex-dependent differences in sensitivity to heat are not well understood in humans, as findings have suggested both greater and lesser susceptibility to heat stroke in women versus men (Giersch et al., 2022). Experimental studies with mice showed females, but not males, exposed to exertional heat stroke exhibited delayed myocardial dysfunction, potentially influencing long-term cardiovascular health (Laitano et al., 2020). Future studies are needed to examine how body size, physiology, cardiovascular health, and energetic expense patterns are linked to temperature fluctuations, hurricane exposures, and mortality across ages in this population.

Our findings complement a body of work examining the effects of early life adversity in humans. Studies have shown that early life adversity is associated with poorer health and reduced longevity in humans (e.g., Barker et al., 2002; Deighton et al., 2018; Gluckman et al., 2008). However, this research faces challenges such as the prevalence of confounding variables like smoking and job insecurity, the difficulty of disentangling effects of different adverse experiences which tend to co-occur, and a reliance on retrospective surveys, which are prone to recall and reporting bias. Animal studies can help overcome these challenges because non-human species are characterized by simpler systems with fewer confounding variables and less clustering of different types of adversity, and non-human animals tend to have shorter lifespans, which allows researchers to observe individuals from birth to death (Dettmer & Chusyd, 2023; Patterson et al., 2023; Snyder-Mackler et al., 2020). Research on early life adversity in other species, especially natural populations, can shed light on the evolutionary pressures shaping early life sensitivities and provide the opportunity to disentangle confounding and correlated environmental factors. In our study, we measured ten forms of adversity and age at death in a large sample of males and females, and were able to show that the form of adversity, socio-sexual context, and other biological factors interact to shape the timing and severity of consequences. Natural populations of non-human animals can prove valuable not only for improving our understanding of the evolutionary pressures that shape developmental plasticity, life history strategies, and early life sensitivities, but also for better contextualizing findings in humans and informing future research in humans.

Our study demonstrates that exposure to early life adversity increases mortality risk in male and female rhesus macaques. Lower odds of surviving to reproductive age indicates that early life adversity can have major fitness ramifications for both an organism and their parents. Reduced life expectancy among those who survive to adulthood, suggests that early life adversity can have persisting fitness costs and long-term health consequences. Adversities related to the maternal social and nutritional environment generally had the largest impacts on offspring survival. We also found sex-dependent effects of early life adversity in this study, which are likely driven by social system characteristics (i.e., female philopatry) and sex-based variation in energetic demands. Further research into the biological mechanisms underlying these survival patterns are needed to better understand how early life adversity impacts fitness and health. Importantly, variation in model estimates and predictions convey that while early life adversity can have negative consequences, such effects are not definitive. Social connections and behavioral adjustments (Campos & Fedigan, 2009; Testard et al., 2021) should be investigated as potential contributors to resilience.

## Acknowledgements

This work was supported by the National Institutes of Health (R01-AG060931; R21AG078554; ORIP-P40OD012217), the National Science Foundation (SMA-2105307), and a collaborative initiative between the National Science Foundation and the European Research Council (supplemental funding for NSF-SMA-2105307 and ERC-864461).

## Supplementary Materials

### Social behavior

Behavioral data were collected continuously using 10-min focal animal samples on handheld computers (Brent, MacLarnon, et al., 2013). From 2010 to 2021, 17 observers conducted focal samples on adults in 6 social groups (groups F, HH, KK, R, S, and V). During focal samples, observers recorded activity state (e.g., resting, traveling, feeding) and social interactions with other adults. For social interactions, observers recorded type of behavior (e.g., grooming, vocal grunts, approaches, contact aggression, threats, displacements), the identity of the partner, and whether the focal or partner initiated the interaction. Behaviors were recorded as instantaneous point occurrences and durational states. Focal animals were selected randomly and data were balanced such that animals were sampled roughly equally across times of day and across the study period. Animals included in our analyses were focal followed for a mean of 5.72 hours per study year (range: 3.17-10.33 hours).

To measure maternal sociality, we calculated a composite sociality index (CSI) using the affiliative social behaviors, approaches and grooming. For each mother in each year, we tabulated the rate of approaches (count of approaches to and from other adult females divided by the number of hours the mother was focalled) and the rate of grooming bouts (count of grooming bouts given to and received from adult females divided by the number of hours the mother was focalled). We calculated the mean approach and grooming rates for all adult females in each social group in each year because these metrics vary across groups and across time. A mother’s approach and grooming rates were divided by the mean rate for a given group-year. These standardized approach and grooming rates were added together and divided by 2 (the number of behaviors) to create the CSI for each mother.

**Table S1.**
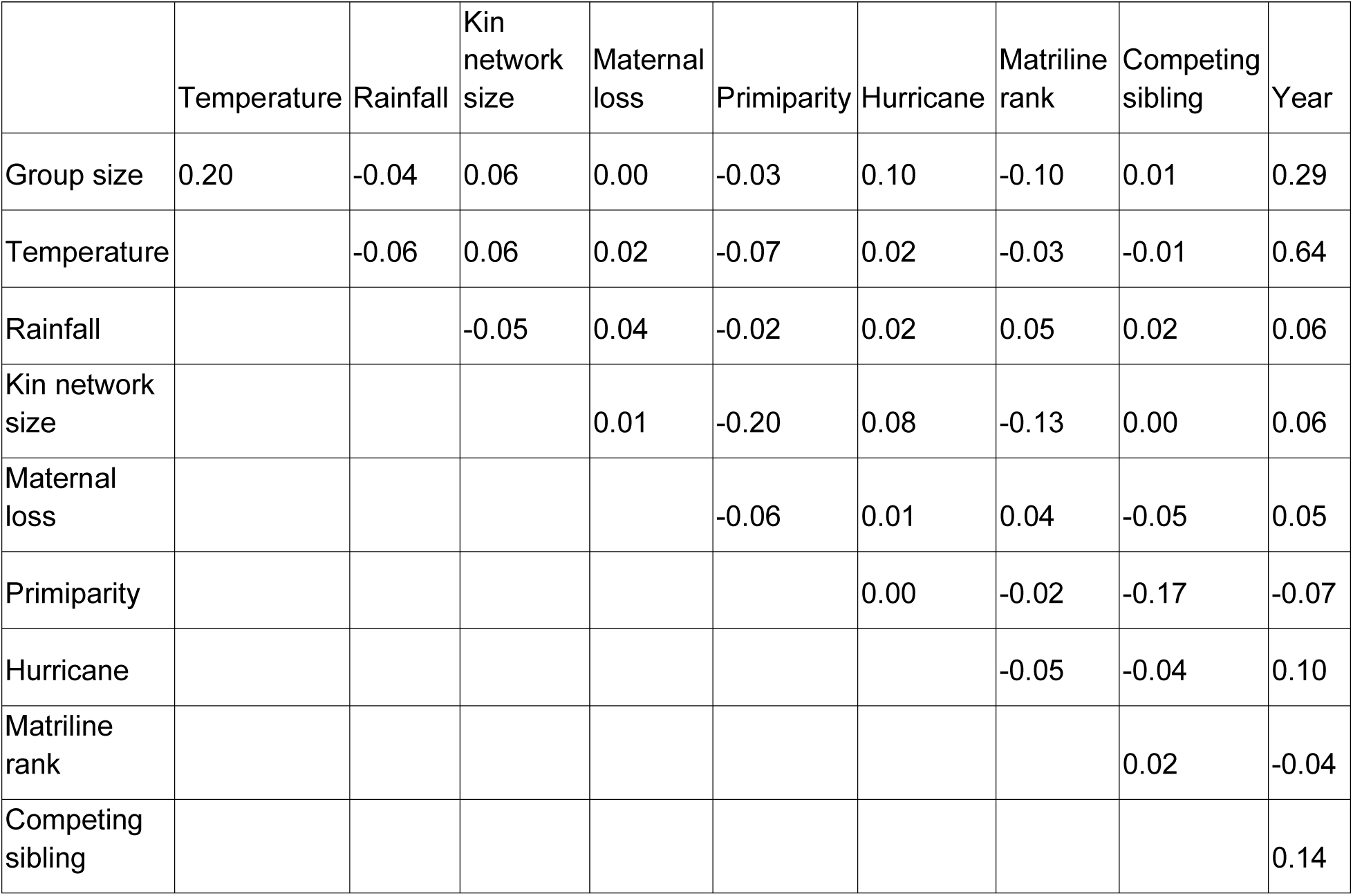
Correlations among model variables.

**Table S2.** Models including and excluding pedigree produce similar predicted effects of early life adversity on survival.

| Early life |  |  |  |  |  |  |  |  |  |
| --- | --- | --- | --- | --- | --- | --- | --- | --- | --- |
| Model including pedigree |  |  |  |  | Model excluding pedigree |  |  |  |  |
|  | Estimate | Est.Error | Q2.5 | Q97.5 |  | Estimate | Est.Error | Q2.5 | Q97.5 |
| Intercept | 2.67 | 0.28 | 2.19 | 3.27 | Intercept | 3.19 | 0.23 | 2.80 | 3.67 |
| ELA Index (continuous, limited) | -0.33 | 0.12 | -0.58 | -0.10 | ELA Index (continuous, limited) | -0.31 | 0.11 | -0.53 | -0.11 |
| Pedigree Intercept | 2.19 | 0.22 | 1.68 | 2.59 |  |  |  |  |  |

| Adults |  |  |  |  |  |  |  |  |  |
| --- | --- | --- | --- | --- | --- | --- | --- | --- | --- |
| Model including pedigree |  |  |  |  | Model excluding pedigree |  |  |  |  |
|  | Estimate | Est.Error | Q2.5 | Q97.5 |  | Estimate | Est.Error | Q2.5 | Q97.5 |
| Intercept | 2.63 | 0.04 | 2.56 | 2.72 | Intercept | 2.63 | 0.04 | 2.57 | 2.71 |
| ELA Index | -0.12 | 0.04 | -0.20 | -0.03 | ELA Index | -0.12 | 0.04 | -0.21 | -0.04 |
| Pedigree Intercept | 0.14 | 0.09 | 0.01 | 0.34 |  |  |  |  |  |

**Table S3.** Models of cumulative early life adversity on survival in early life and adulthood.

| Early life |  |  |  |  | Adults |  |  |  |  |
| --- | --- | --- | --- | --- | --- | --- | --- | --- | --- |
|  | Estimate | Est.Error | Q2.5 | Q97.5 |  | Estimate | Est.Error | Q2.5 | Q97.5 |
| Intercept | 1.03 | 0.05 | 0.93 | 1.12 | Intercept | 2.77 | 0.02 | 2.73 | 2.80 |
| Sex (male) | -0.05 | 0.10 | -0.24 | 0.15 | Sex (male) | -0.13 | 0.02 | -0.18 | -0.08 |
| Cumulative early life adversity index | -0.29 | 0.07 | -0.43 | -0.15 | Cumulative early life adversity index | -0.04 | 0.02 | -0.08 | -0.01 |
| Sex*Cumulative early life adversity index | -0.07 | 0.09 | -0.24 | 0.12 | Sex*Cumulative early life adversity index | 0.00 | 0.02 | -0.04 | 0.05 |
| Birth year intercept | 3.37 | 0.33 | 2.78 | 4.04 | Birth year intercept | 0.03 | 0.02 | 0.00 | 0.07 |
| Maternal ID intercept | 0.78 | 0.09 | 0.60 | 0.96 | Maternal ID intercept | 0.07 | 0.03 | 0.01 | 0.13 |

**Table S4.**
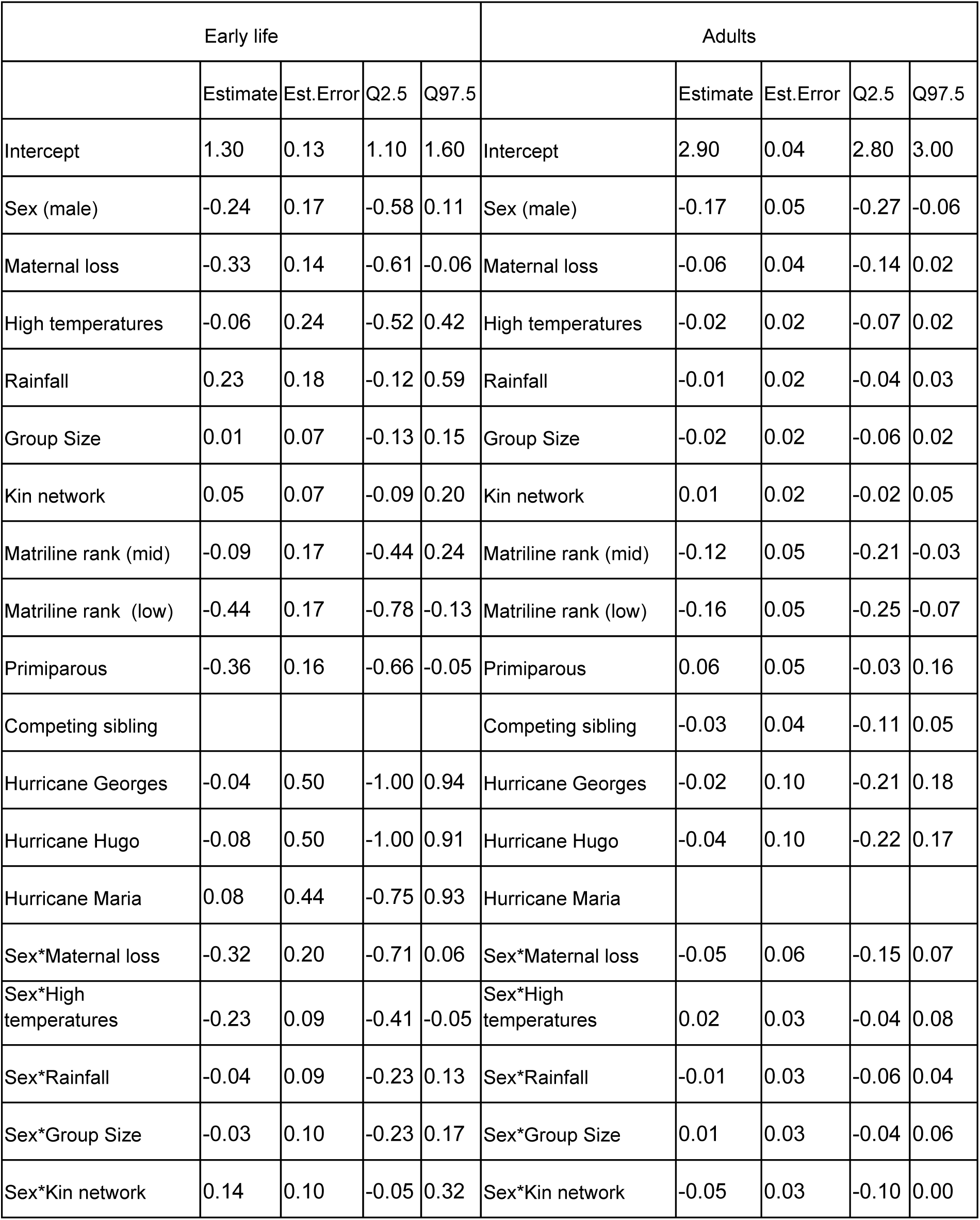
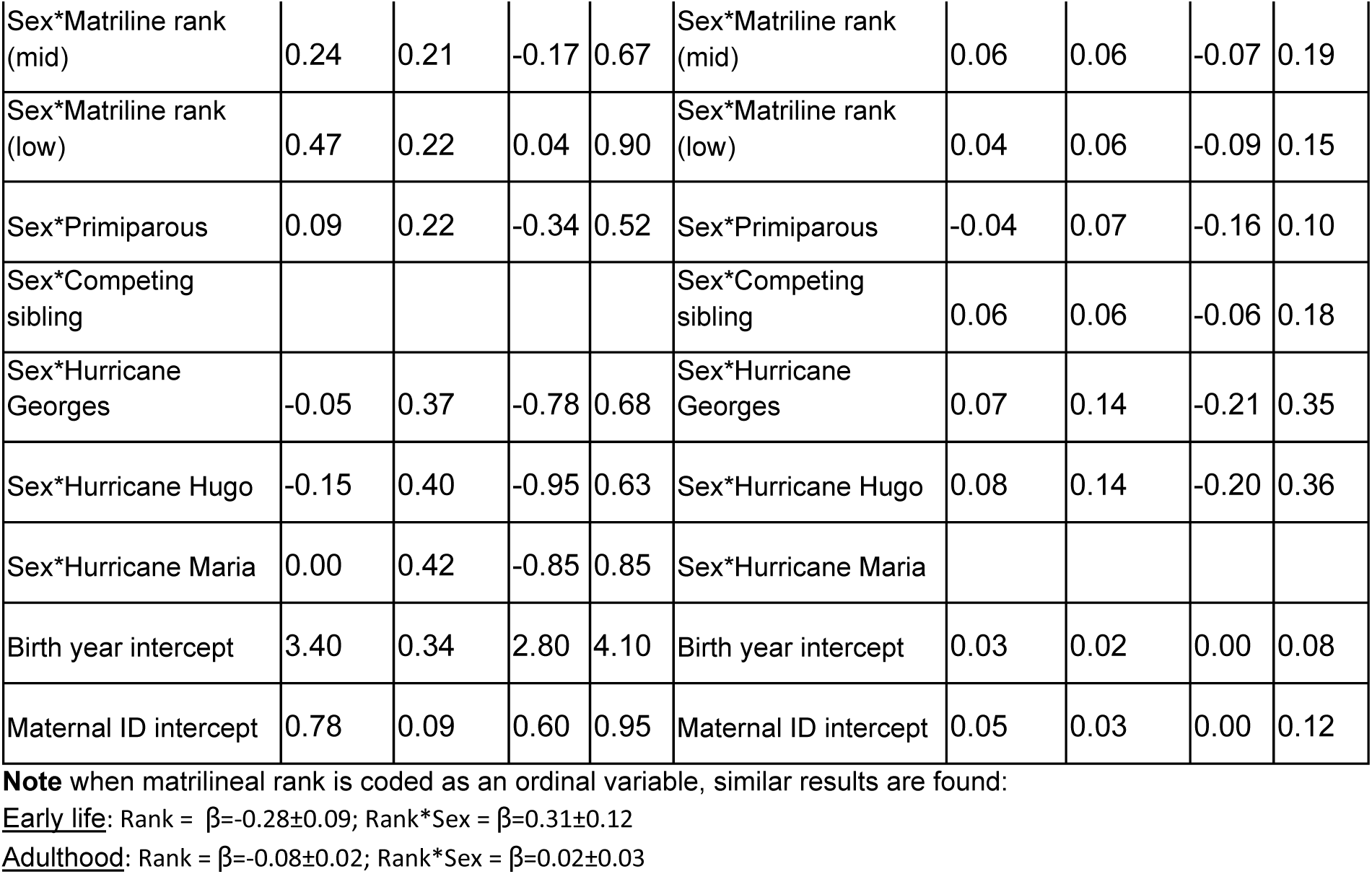
Survival as a function of individual forms of early life adversity.

| Early life |  |  |  |  | Adults |  |  |  |  |
| --- | --- | --- | --- | --- | --- | --- | --- | --- | --- |
|  | Estimate | Est.Error | Q2.5 | Q97.5 |  | Estimate | Est.Error | Q2.5 | Q97.5 |
| Intercept | 1.30 | 0.13 | 1.10 | 1.60 | Intercept | 2.90 | 0.04 | 2.80 | 3.00 |
| Sex (male) | -0.24 | 0.17 | -0.58 | 0.11 | Sex (male) | -0.17 | 0.05 | -0.27 | -0.06 |
| Maternal loss | -0.33 | 0.14 | -0.61 | -0.06 | Maternal loss | -0.06 | 0.04 | -0.14 | 0.02 |
| High temperatures | -0.06 | 0.24 | -0.52 | 0.42 | High temperatures | -0.02 | 0.02 | -0.07 | 0.02 |
| Rainfall | 0.23 | 0.18 | -0.12 | 0.59 | Rainfall | -0.01 | 0.02 | -0.04 | 0.03 |
| Group Size | 0.01 | 0.07 | -0.13 | 0.15 | Group Size | -0.02 | 0.02 | -0.06 | 0.02 |
| Kin network | 0.05 | 0.07 | -0.09 | 0.20 | Kin network | 0.01 | 0.02 | -0.02 | 0.05 |
| Matriline rank (mid) | -0.09 | 0.17 | -0.44 | 0.24 | Matriline rank (mid) | -0.12 | 0.05 | -0.21 | -0.03 |
| Matriline rank (low) | -0.44 | 0.17 | -0.78 | -0.13 | Matriline rank (low) | -0.16 | 0.05 | -0.25 | -0.07 |
| Primiparous | -0.36 | 0.16 | -0.66 | -0.05 | Primiparous | 0.06 | 0.05 | -0.03 | 0.16 |
| Competing sibling |  |  |  |  | Competing sibling | -0.03 | 0.04 | -0.11 | 0.05 |
| Hurricane Georges | -0.04 | 0.50 | -1.00 | 0.94 | Hurricane Georges | -0.02 | 0.10 | -0.21 | 0.18 |
| Hurricane Hugo | -0.08 | 0.50 | -1.00 | 0.91 | Hurricane Hugo | -0.04 | 0.10 | -0.22 | 0.17 |
| Hurricane Maria | 0.08 | 0.44 | -0.75 | 0.93 | Hurricane Maria |  |  |  |  |
| Sex*Maternal loss | -0.32 | 0.20 | -0.71 | 0.06 | Sex*Maternal loss | -0.05 | 0.06 | -0.15 | 0.07 |
| Sex*High temperatures | -0.23 | 0.09 | -0.41 | -0.05 | Sex*High temperatures | 0.02 | 0.03 | -0.04 | 0.08 |
| Sex*Rainfall | -0.04 | 0.09 | -0.23 | 0.13 | Sex*Rainfall | -0.01 | 0.03 | -0.06 | 0.04 |
| Sex*Group Size | -0.03 | 0.10 | -0.23 | 0.17 | Sex*Group Size | 0.01 | 0.03 | -0.04 | 0.06 |
| Sex*Kin network | 0.14 | 0.10 | -0.05 | 0.32 | Sex*Kin network | -0.05 | 0.03 | -0.10 | 0.00 |
| Sex*Matriline rank (mid) | 0.24 | 0.21 | -0.17 | 0.67 | Sex*Matriline rank (mid) | 0.06 | 0.06 | -0.07 | 0.19 |
| Sex*Matriline rank (low) | 0.47 | 0.22 | 0.04 | 0.90 | Sex*Matriline rank (low) | 0.04 | 0.06 | -0.09 | 0.15 |
| Sex*Primiparous | 0.09 | 0.22 | -0.34 | 0.52 | Sex*Primiparous | -0.04 | 0.07 | -0.16 | 0.10 |
| Sex*Competing sibling |  |  |  |  | Sex*Competing sibling | 0.06 | 0.06 | -0.06 | 0.18 |
| Sex*Hurricane Georges | -0.05 | 0.37 | -0.78 | 0.68 | Sex*Hurricane Georges | 0.07 | 0.14 | -0.21 | 0.35 |
| Sex*Hurricane Hugo | -0.15 | 0.40 | -0.95 | 0.63 | Sex*Hurricane Hugo | 0.08 | 0.14 | -0.20 | 0.36 |
| Sex*Hurricane Maria | 0.00 | 0.42 | -0.85 | 0.85 | Sex*Hurricane Maria |  |  |  |  |
| Birth year intercept | 3.40 | 0.34 | 2.80 | 4.10 | Birth year intercept | 0.03 | 0.02 | 0.00 | 0.08 |
| Maternal ID intercept | 0.78 | 0.09 | 0.60 | 0.95 | Maternal ID intercept | 0.05 | 0.03 | 0.00 | 0.12 |

**Note** when matrilineal rank is coded as an ordinal variable, similar results are found:
Early life: Rank = $\beta = -0.28 \pm 0.09$ ; Rank\*Sex = $\beta = 0.31 \pm 0.12$
Adulthood: Rank = $\beta = -0.08 \pm 0.02$ ; Rank\*Sex = $\beta = 0.02 \pm 0.03$

**Table S5.** Maternal social connectedness and offspring survival in early life and adulthood.

| Early life |  |  |  |  | Adults |  |  |  |  |
| --- | --- | --- | --- | --- | --- | --- | --- | --- | --- |
|  | Estimate | Est. Error | Q2.5 | Q97.5 |  | Estimate | Est. Error | Q2.5 | Q97.5 |
| Intercept | 0.93 | 0.16 | 0.62 | 1.25 | Intercept | 2.38 | 0.25 | 1.99 | 2.97 |
| Sex (male) | 0.22 | 0.24 | -0.26 | 0.70 | Sex (male) | 0.15 | 0.19 | -0.21 | 0.56 |
| Maternal social connectedness | 0.15 | 0.17 | -0.19 | 0.49 | Maternal social connectedness | -0.02 | 0.12 | -0.26 | 0.24 |
| Sex*Maternal social connectedness | 0.29 | 0.25 | -0.19 | 0.81 | Sex*Maternal social connectedness | -0.00 | 0.19 | -0.40 | 0.38 |
| Birth year intercept | 2.18 | 0.72 | 1.21 | 3.94 | Birth year intercept | 0.20 | 0.17 | 0.01 | 0.63 |
| Maternal ID intercept | 0.66 | 0.34 | 0.04 | 1.30 | Maternal ID intercept | 0.20 | 0.16 | 0.01 | 0.59 |

**Table S6.** Model comparisons using the loo_compare function in the “brms” R package.

| Early life survival |  |  | Adult survival |  |  |
| --- | --- | --- | --- | --- | --- |
| Model | elpd_diff | se_diff | Model | elpd_diff | se_diff |
| Index | 0.0 | 0.0 | Index | 0.0 | 0.0 |
| Multivariate | -1.0 | 4.4 | Multivariate | -3.3 | 5.9 |

**Figure S1.**
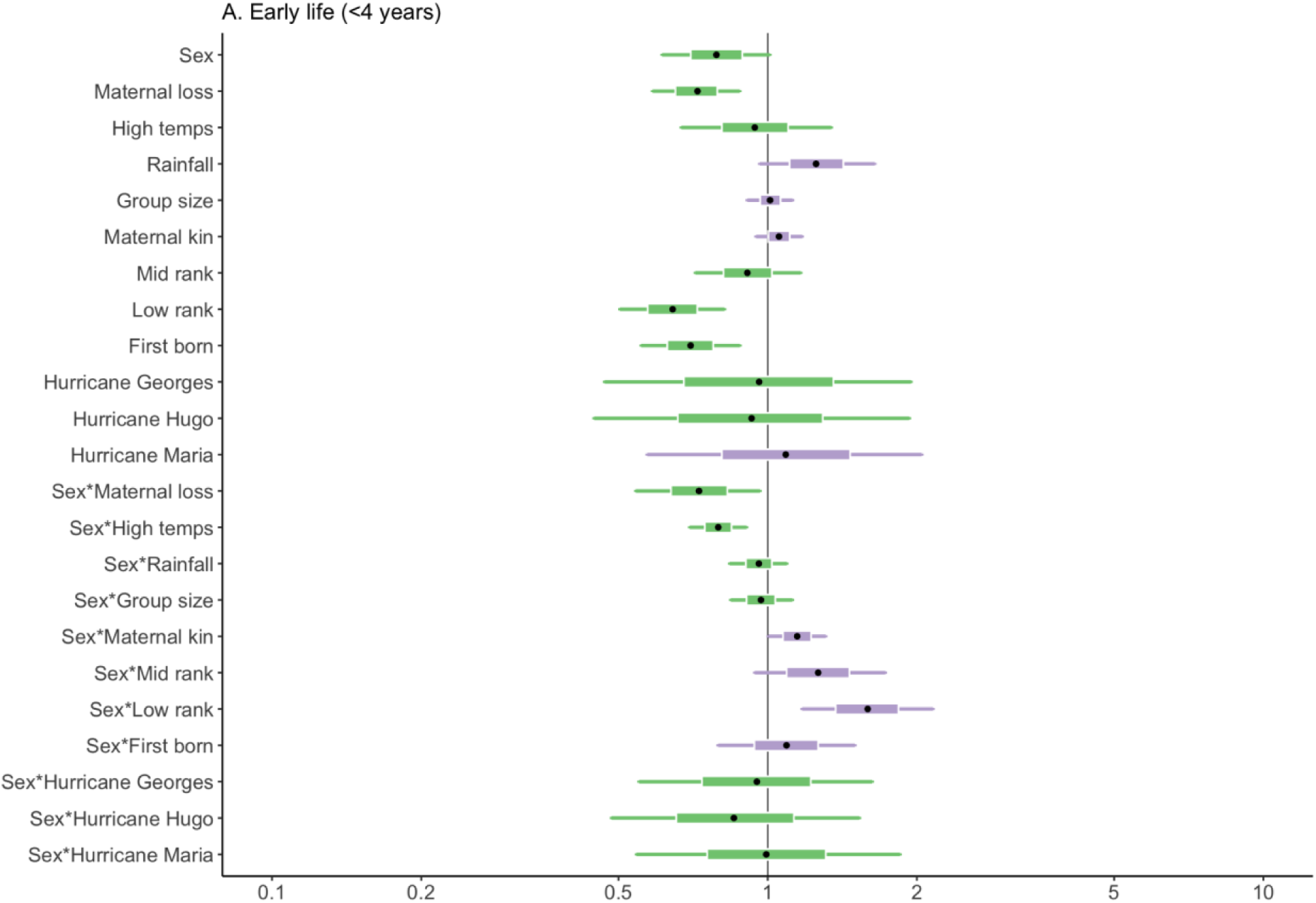
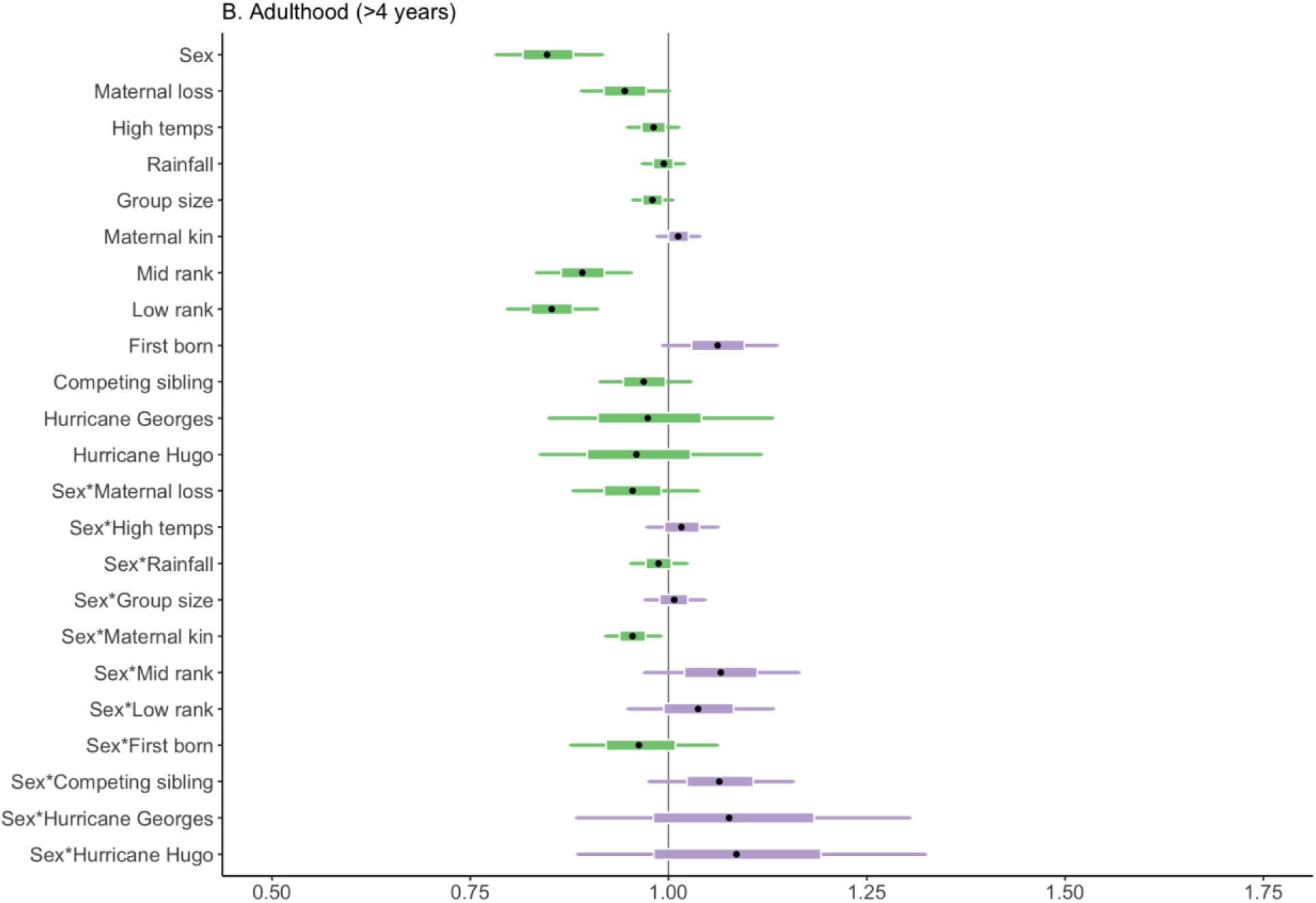
Model effects of sex and the forms of early life adversity on survival during early life (A) and adulthood (B). The outer bars show the 85% credible intervals, the inner boxes show the 50% credible intervals, and the black circles in the middle show the medians of the posterior distributions. Green shading represents negative effect sizes and purple shading represents positive effect sizes.

## Notes

https://github.com/skpatter/ELA_Survival_Macaques

### Competing Interest Statement

The authors have declared no competing interest.

